# Unraveling Fitness Landscapes in Plant-Associated Bacteroidota

**DOI:** 10.64898/2025.12.01.691665

**Authors:** Marta Torres, Vincent Lombard, Nicolas Terrapon, Adam M. Deutschbauer

## Abstract

Members of the bacterial phylum Bacteroidota inhabit a wide range of ecosystems. Although significant progress has been made over the past decade in understanding the metabolic functions of clinically relevant *Bacteroidota*, far less is known about the environmental species. Despite being one of the dominant phyla linked to crops, there is limited information on the mechanisms that enable Bacteroidota to thrive in plant and soil environments. A key obstacle in understanding gene fitness in Bacteroidota is the lack of genome-wide functional studies, largely due to their inherent resistance to antibiotics, which complicates genetic manipulation. In this study, we used randomly barcoded transposon mutagenesis sequencing (RB-TnSeq) to measure gene fitness in a plant-associated Bacteroidota, *Mucilaginibacter yixingensis* YX-36. Our data sheds light on pathways involved in rhizosphere colonization, gliding motility, stress tolerance, and carbon metabolism. Notably, we found that phylum-specific genes such as Polysaccharide Utilization Loci (PULs) and Carbohydrate-Active enzymes (CAZymes) are needed for fitness in the plant niche. Overall, this work advances our understanding of gene functions in environmental Bacteroidota species and provides a foundation for future research on their roles in plant–microbe interactions.

## 1. Introduction

Members of the phylum Bacteroidota colonize a wide range of ecological niches, including water (ocean, freshwater), soil, plants, and animals. In the latter, they can be found on the skin, in the oral cavity, and in the gastrointestinal tract, where they account for a major fraction of the bacteriome and play key roles in various biological functions, such as nutrient supply, immune system regulation, and disease protection[1]. Although they are best studied as dominant members of the gut microbiota[2], their roles in other environments remain poorly characterized, and less attention has been given to Bacteroidota in plant microbiomes compared to their counterparts associated with humans and animals[3].

In the environment, a major source of renewable carbon and energy for microbial growth comes from plants. Plants are notably rich in glycans that range from simple compounds (mono-/disaccharides) to complex structures (polysaccharides), the latter being more recalcitrant to degradation and therefore more persistent in soil[4]. Plant glycans comprise a major fraction of plant biomass and root mucilage, and they have extensively been studied in different plant models, such as in *Brachypodium distachyon*[5, 6], in which the cell wall is predominantly composed of arabinoxylans, cellulose, and mixed linkage glucans [(1,3;1,4)-β-D-glucans][7]. While mycorrhizal and saprophytic fungi were once considered the main degraders of plant polysaccharides, soil bacteria such as members from the phylum Bacteroidota are now recognized as key contributors[8, 9].

Bacteroidota have been detected in plant and soil samples from various locations, including agriculture fields[10] and unmanaged forests[11]. They can account for a large fraction of the root microbiome[12, 13] and they have also been detected in leaves, for instance of gymnosperm trees[14, 15]. In some angiosperm species, they are the main phylum enriched in roots and rhizosphere *versus* bulk soil[16]. Notably, they tend to be more common in the roots and rhizosphere of wild crops than in domesticated plants[17–19]. In recent years, Bacteroidota have been proposed as an ecologically important player in soil functioning and are therefore recommended as a biological indicator of agricultural soil health[20, 21], as they are significantly affected by agricultural practices. In addition, they are potential candidates to be used as microbial inoculants to replace traditional fertilizers. For instance, several *Mucilaginibacter* and *Flavobacterium* strains have shown promising effects on plant growth and alleviation of biotic and abiotic stresses[22–24].

The wide distribution of Bacteroidota members and their diverse lifestyles suggest high adaptability to various environments. This ubiquity exemplifies their impressive versatility in niche adaptation and genomic plasticity. Living in complex habitats and metabolizing many different substrates, environmental species tend to have larger genomes than non-environmental species, which can be correlated with their broad catabolic capabilities[1]. The Bacteroidota are well-known specialists for the degradation of high molecular weight organic matter, especially in the form of polysaccharides and proteins[25]. They thrive thanks to their ability to secrete diverse arrays of carbohydrate-active enzymes (CAZymes; glycoside hydrolases and polysaccharide lyases that break down glycosidic bonds, and carbohydrate binding modules that bind to carbohydrates and direct the catalytic machinery towards them[26]) that target the highly varied and complex glycans (e.g. pectins, hemicelluloses like xylans) found in the soil and plant environment[4, 25]. Interestingly, among environmental members, terrestrial Bacteroidota show a significantly higher abundance and diversity of CAZymes[3], which reflects that terrestrial Bacteroidota are highly adapted to plant carbohydrate metabolism, which in turn appears to be critical for their profusion in the plant environment[9, 27].

Efficient degradation of complex polysaccharides requires tight coordination between enzymes and transport systems. Bacteroidota achieve this through an energy-saving system of genomic organization, whereby collaborating CAZymes, transporters, and regulators are grouped into Polysaccharide Utilization Loci (PULs). These loci enable high level production of specific CAZymes only when their substrate glycans are abundant in the local environment[25]. All this can be further enhanced by the phylum-specific Type IX Secretion System (T9SS), which is highly effective at secreting CAZymes and/or tethering them to the cell surface, and is tightly coupled to the ability to rapidly glide over solid surfaces, a connection that promotes an active hunt for nutrition[4, 28]. This gives the Bacteroidota an advantage over other species in a competitive environment as they can sequester oligosaccharides away from the competition, and degrade them in their periplasm thanks to the synergy of PUL systems, while other species can only degrade them outside of the cells and share them[29]. Moreover, Bacteroidota play a pivotal role in community functioning by breaking down complex plant polysaccharides and contributing to cross-feeding networks within soil microbial communities[30, 31]. This results in a blow to their own fitness for supplying others, as it happens with other microbes that degrade special food/polymers and produce important secretions for use in a community[32].

Determining the mechanisms that enable Bacteroidota to colonize plant roots may provide opportunities for enhancing crop production through microbiome engineering. However, despite the interest in unravelling the functional capabilities of environmental Bacteroidota, it remains challenging to infer the pathways and mechanisms involved in such phenotypes. Great advances in understanding biological processes result from determining the function of individual genes. One approach to identify the underlying genetic mechanisms involved in bacterial growth in a particular condition is to disrupt genomic regions and then screen the mutant library for a given phenotype. Among the most scalable high-throughput techniques is randomly barcoded transposon mutagenesis sequencing (RB-TnSeq)[33], that combines the advantages of TnSeq[34] and rapid quantification of each transposon mutant using unique DNA barcodes. This approach has successfully been conducted in dozens of bacteria[35], most of them belonging to the phylum Pseudomonadota and only a few to the phylum Bacteroidota[36]. This unbalance partly reflects the more challenging genetic manipulation of Bacteroidota species. Examples of Bacteroidota in which TnSeq has been applied for fitness evaluation are: *Echinicola vietnamensis*[37], *Pedobacter* sp.[37], *Pontibacter actiniarum*[37], *Porphyromonas gingivalis*[38]*, Bacteroides thetaiotaomicron*[39–41], *B. ovatus*[42], *B. cellulosilyticus*[42] and *B. fragilis*[43]. These libraries have mainly been assayed for metabolism of carbon compounds and animal gut colonization.

To address this knowledge gap and assess the genetic fitness determinants of plant colonization in environmental Bacteroidota, we attempted the construction of RB-TnSeq mutant libraries in representatives of various soil and plant-related genera, including *Flavitalea*, *Flavisolibacter*, *Niastella*, *Mucilaginibacter*, *Chitinophaga*, *Runella*, *Sediminibacterium*, *Spirosoma*, and *Dyadobacter*. Due to the inherent antibiotic resistance of many strains, we only succeeded with *M. yixingensis* YX-36, a species belonging to the family Sphingobacteriaceae that was originally isolated from a vegetable soil[44]. Members of the genus *Mucilaginibacter* are among the most abundant genera found in agricultural soils[45], and several species have been shown to promote plant growth[22, 46, 47]. Because of their relevance, *Mucilaginibacter* isolates are frequently included in synthetic communities (SynComs) aimed at studying bacterial interactions with plant models, like *B. distachyon*[48, 49]. However, the genetic traits underlying the ability of *Mucilaginibacter* species to successfully colonize the plant rhizosphere remain largely unexplored. Our findings indicate that gliding motility and the degradation of complex carbon sources are key factors for effective rhizosphere colonization by *M. yixingensis*. Furthermore, the tools and insights presented here will serve as valuable resources for further investigations into gene function within environmental Bacteroidota.

## 2. Material and Methods

### 2.1 Bacterial strains and culture conditions

*Escherichia coli* strains were cultured in LB at 37°C. The environmental Bacteroidota strains (**Table 1**), either obtained from the DSMZ German strain collection https://www.dsmz.de/ or originally isolated from a switchgrass (*Panicum virgatum*) field in Oklahoma (United States), were routinely cultured in R2A at 30°C. If needed, 180 rpm rotary shaking was used. Antibiotic susceptibility of the Bacteroidota strains was conducted in R2A supplemented with different antibiotics: tetracycline (Tc), chloramphenicol (Cm), streptomycin (Str), kanamycin (Km), erythromycin (Erm) and gentamicin (Gm) used at a final concentration of 50 and/or 100 µg/mL. When appropriate, diaminopimelic acid (DAP) was added to a final concentration of 300 µM, and phenylalanine-arginine-β-naphthylamide (PAβN) was used at 25 µg/mL.

**Table 1.**
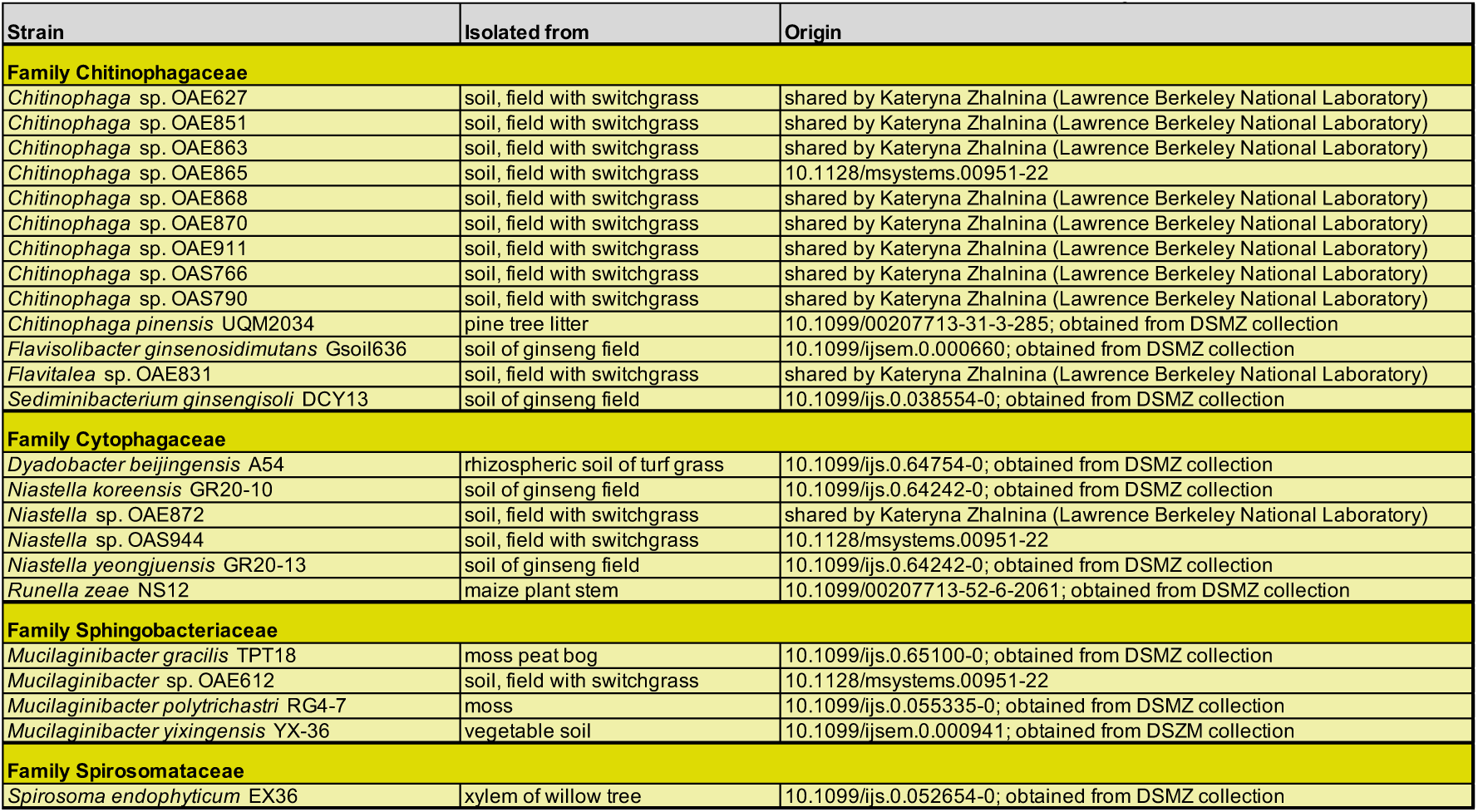
Bacteroidota environmental strains used in this study.

### 2.2 Construction of barcoded mutant libraries in environmental Bacteroidota

#### a) Identification of an optimal vector for transposon mutagenesis using the magic pool approach

This methodology is a combinatorial strategy for constructing a catalog of thousands of uniquely assembled transposon delivery vectors from a collection of DNA parts (transposases, promoters, and selection markers)[37]. It streamlines the process of identifying vectors that have the optimal combination of parts by simultaneously testing the efficacy of all possible vectors in the catalog to generate mutants in a new species of interest[50]. Once a vector construction (i.e. the different parts) is identified as the most efficient, it can be reassembled and used to generate a final mutant library. The erythromycin magic pool used in this study (AMD1241) was previously described by our lab[37]. AMD1241 is a mix of Tn5 (AMD279) and mariner (AMD280) magic pools. In the Tn5-Erm magic pool, there is part1 (Tn5 transposase), ten variants of the antibiotic resistance gene upstream region (part2), five variants of the Erm antibiotic resistance gene (part3), 25 variants of the transposase upstream region (part5) and a randomly barcoded part4, resulting in 1,250 possible vector combinations[37]. For the mariner-Erm magic pool, part1 is a mariner transposase, and there are the same ten variants of part2 and the same five variants of part3, and 24 variants of part5, giving 1,200 possible vector combinations[37]. DNA part sequences and sources are available from Liu et al[37].

Conjugations between the Erm-sensitive Bacteroidota strains and the Erm magic pool (AMD1241) were conducted in both LB and R2A supplemented with DAP using conjugation disks. After 16 h, the conjugation was scraped off, resuspended in LB or R2A, and plated in selective media with the Erm, with and without PAβN. Plates were incubated at 30°C for 48 h to let visible colonies develop. When possible, preliminary libraries were constructed by pooling around 1,000 colony forming units (CFU) and growing them for two population doublings in liquid media. We then performed TnSeq to characterize each mutant library by mapping the insertion location and its associated random DNA barcode as previously described[33], thereby simultaneously assessing the efficacy of the vectors in the magic pool. To map the genomic locations of the transposon insertions and link these insertions to their associated DNA barcodes, we used a variation of previously described TnSeq protocol[33], where we use two rounds of PCR to selectively enrich for transposon junctions[51].

#### b) Construction of a final mutant library with the optimal transposon delivery vector

Once a vector design in the magic pool was identified as the most effective (i.e. most prevalent and least biased and hence most successful at mutant generation), the single reassembled vector with the optimal combination of parts was used to construct a final RB-TnSeq[33] transposon mutant library. We only identified an efficient vector assembly for *M. yixingensis* YX-36, a strain originally isolated from a vegetable soil [44]. The vector used was pTGG43_NN2[37]. Conjugation between YX-36 and an *E. coli* conjugation donor carrying pTGG43_NN2 was performed as explained above. Plates were incubated at 30°C for 48 h. We then pooled thousands of colonies and grew the library in liquid R2A supplemented with Erm, added glycerol to a final volume of 15%, made multiple 1-mL −80°C freezer stocks (∼10^8^ cells/mL) of the final library for subsequent experiments, and collected cell pellets to extract genomic DNA for TnSeq mapping. To map the genomic locations of the transposon insertions and link these insertions to their associated DNA barcodes, we used the TnSeq protocol[33] with two rounds of PCR as explained above. The final library was called *Mucilaginibacter*_YX36_ML5.

### 2.3 Fitness assays with *Mucilaginibacter*_YX36_ML5

Plant and *in vitro* conditions were chosen according to traits previously described in *Mucilaginibacter* species and in general, other environmental Bacteroidota species. These include colonization of the rhizosphere of *Brachypodium* plants[48, 49]; degradation of polysaccharides[9, 52, 53]; and tolerance to phosphate[54] and metals[55].

The different plant and *in vitro* fitness assays were conducted with *Mucilaginibacter*_YX36_ML5 transposon library as follows. Every time experiments were conducted, one aliquot of the glycerol stocks containing the *Mucilaginibacter*_YX36_ML5 transposon library was thawed and inoculated in 25 mL fresh R2A with 50 µg/mL Erm and grown for approximately 16 h at 30°C and 180-rpm shaking until the culture reached OD600 ∼1.3. Time0 samples enumerating the relative abundance of each mutant were collected by centrifuging 1 mL aliquots and frozen at -20°C until further DNA purification. The remaining cells were then washed twice with a chemically-defined media without a carbon source (RCH2_defined_noCarbon)[56] prior to inoculation. A summary of the 85 genome-wide fitness assays performed in this study with the *Mucilaginibacter*_YX36_ML5 mutant library, which include *in vitro* and plant assays, as well as the number of replicates used per condition, can be found in **Table S1**.

#### a) *In vitro* fitness assays

Assays were conducted in defined media or rich media depending on the type of experiment, at starting OD600=0.03. Carbon assays were conducted in RCH2_defined_noCarbon[56] supplemented with different carbon sources (i.e. monosaccharides, disaccharides, polysaccharides). After 24 or 48 h, cultures with an OD higher than the average non-inoculated controls plus two times the standard deviation were centrifuged. Stress assays were conducted in R2A supplemented with compounds at inhibitory concentrations. Temperature assays were conducted in R2A. After 24, 48, 72 h or 96 h, cultures with an OD lower than the no stress controls were centrifuged. Pellets were kept frozen at -20°C until further DNA purification.

For gliding motility assays, the mutant library was inoculated into the center of 0.5% w/v agar R2A plates, and ‘outer’ samples with motile cells were removed with a sterile razor after 24 h[56]. Cells were harvested from agar by centrifugation, and pellets were kept at -20°C until DNA purification.

#### b) Plant colonization fitness assays

*Brachypodium distachyon* Bd21-3 seeds were dehusked and sterilized in 70% v/v ethanol for 30 s, and in 6% v/v NaOCl for 5 min, followed by five wash steps in sterile water. Seeds were stratified in the dark for 2 to 3 days at 4°C. After stratification, seeds were germinated on 1 % (w/v) water-agar plates in a 130 µmol/m^2^ s^−1^ 16-h light/8-h dark regime at 24°C for 3 days. Then, using sterile forceps, seedlings were transferred to Magenta GA-7 plant culture boxes filled with ∼ 150 mL of 0.5X Murashige & Skoog (MS)[57] basal salts (MSP01, Caisson Laboratories, United States) with 0.3% (w/v) agar. Boxes were covered with vented lids and maintained in the same conditions as explained above. Fifteen days after germination, 1 mL of *Mucilaginibacter*_YX36_ML5 washed cells (OD600=1) were diluted in 5 mL of 0.5X MS and the whole volume was inoculated into the plants by injecting directly on the MS agar substrate in 6-7 different spots at ∼ 1 cm around the plant stem. The substrate was stirred with a sterile spatula around the plant to homogenize as much as possible the bacterial inoculum. Two harvest points were considered, 7 and 21 days post inoculation (dpi). Briefly, plants were uprooted, and the aerial part (root/shoot junction) was cut using sterile scissors and disposed of. Isolated roots were collected in 50-mL falcons containing 20 mL of R2A and vortexed at high speed for 2 minutes; this sample was referred to as the ‘rhizosphere’. Samples were then incubated (outgrown) for 16 h at 30°C and 180-rpm shaking in R2A. Two-mL samples from all cultures were harvested. After centrifugation (3,000 g for 3 min), pellets were stored at - 20°C until DNA purification.

### 2.4 DNA isolation, library preparation, sequencing and fitness value calculation

DNA from frozen pellets was isolated using the DNeasy Blood & Tissue Kit (Qiagen, Germany) according to manufacturer’s instructions. Purified RNA-free DNA was used as a template for DNA barcode PCR amplification using previously described PCR conditions[33]. Strain and gene fitness calculations were done using the computation pipeline developed by Wetmore et al.[33]; code available at bitbucket.org/berkeleylab/feba/src/master. Fitness values for each gene were calculated as the log2 of the ratio of relative barcode abundance for mutants in that gene following library growth in a given condition (i.e. plant colonization) divided by relative abundance in the Time0 (input) sample. A t-like statistic[33] was also determined for every gene in each experiment, which takes into account the consistency of the fitness of all the mutants of that gene to assess if the fitness value is reliably different from 0. To avoid considering weak colonization fitness values, which may not be biologically meaningful, we focused on genes that had |fitness| ≥ 1, with an associated |t| ≥ 3.

### 2.5 Data analysis and data availability

Analysis of gene fitness values was done in R[58]. Graphs and heatmaps were plotted in R using the *ggplot2* package[59]. Raw sequencing data is available on the Figshare website (https://doi.org/10.6084/m9.figshare.30731249.v1). The majority of the fitness data from this study is publicly available at the Fitness Browser website (https://fit.genomics.lbl.gov)[56] for comparative analysis. Experiments that passed and failed our thresholds for inclusion in the Fitness Browser are noted in **Table S2**. The CAZy database (www.cazy.org[26]) was used to annotate CAZy domains. Such annotation results from a semi-manual process, with automated annotation only for sequences reaching significant coverage and similarity to previously annotated full-length sequences, while sequences with low or partial similarity are curated through human expertise. The PUL database (PULDB)[60] (www.cazy.org/PULDB/) was used to identify PULs.

## 3. Results and discussion

### 3.1 Erythromycin and gentamicin are the best candidates for antibiotic selection of environmental Bacteroidota

To determine which antibiotics could be used for selecting mutants in environmental Bacteroidota, we systematically tested the sensitivity of 24 strains to 6 antibiotics commonly used in bacterial genetics. We found that the environmental Bacteroidota strains tested are naturally resistant to most antibiotics, including tetracycline, chloramphenicol, streptomycin and kanamycin. Only erythromycin (Erm) and gentamicin (Gm) were effective on ∼50% of the strains (**Fig. 1**), albeit high concentrations (100 µg/mL) were required for some strains. Given our past success with erythromycin selection as a marker for making RB-TnSeq libraries, we focused our efforts on the Erm-sensitive strains. We tested the combined *mariner* and Tn5 magic pool[37] with Erm selection (AMD1241) on strains *Chitinophaga* sp. OAE865, *Chitinophaga* sp. OAE868, *Niastella* sp. OAE872, *Niastella* sp. OAS944, *Flavitalea* sp. OAE831, *Mucilaginibacter* sp. OAE612, *Mucilaginibacter gracilis* TPT18, *Mucilaginibacter yixingensis* YX-36, *Mucilaginibacter polytrichastri* RG4-7, *Runella zeae* NS12, *Spirosoma endophyticum* EX36, and *Sediminibacterium ginsengisoli* DCY13 (**Fig. 1**). Despite trying different approaches (e.g. different conjugation media, type of transposon delivery vector, selection with different antibiotic concentration, and use of the efflux inhibitor PAβN to reduce antibiotic resistance[61, 62]), we only succeeded in generating transposon mutants in strain YX-36 (**Fig. S1**). Generation of antibiotic-resistant colonies requires that a number of independent events occur efficiently[63]: 1) the DNA must be transferred into the cells, 2) the DNA must not be immediately degraded by host defense systems, 3) the antibiotic resistance genes must be expressed and they must confer antibiotic resistance on the cells and 4) the resistance elements must be maintained as the bacteria multiply, which requires either independent replication of the plasmid or integration of the plasmid (or transposable element) into the host genome. Our results on low numbers of real transconjugants point to a lack of stable plasmid replication, but we cannot rule out other limiting events in the tested environmental strains.

**Figure 1.**
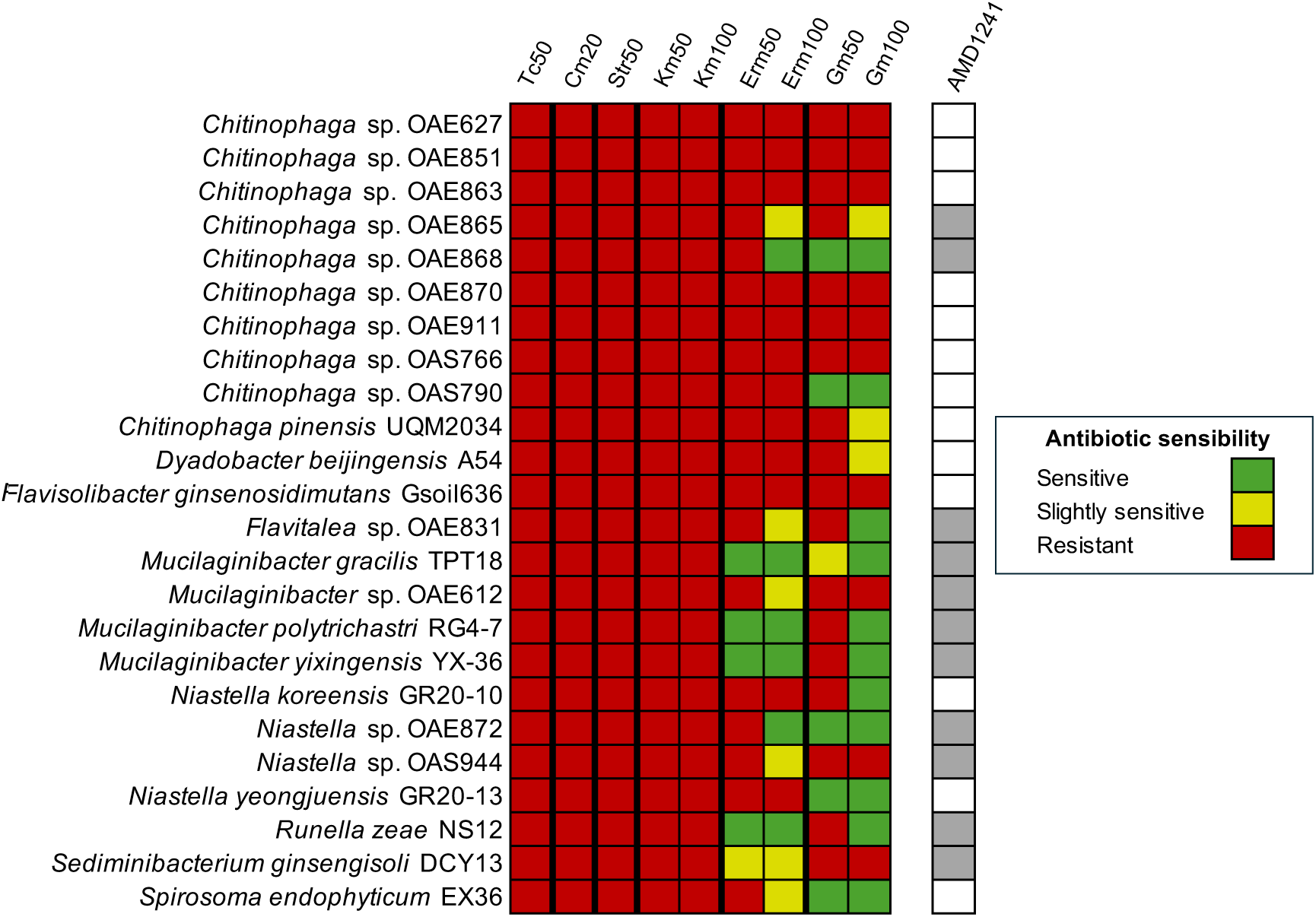
Sensitivity of environmental Bacteroidota strains to tetracycline (Tc), chloramphenicol (Cm), streptomycin (Str), kanamycin (Km), erythromycin (Erm) and gentamicin (Gm). Erm-sensitive strains in which the combined *mariner* and Tn5 magic pool (AMD1241) was used are highlighted. Details on the listed strains can be found in Table 1.

### 3.2 Construction of a barcoded mutant library in *M. yixingensis* YX-36

Despite the widespread antibiotic resistance of the environmental strains tested and the challenge of constructing mutant libraries, we got promising results for one of the strains, *M. yixingensis* YX-36 (NCBI taxonomy ID: 1295612) (**Fig. S1**). Although a draft genome assembly was already available (NCBI RefSeq assembly GCF_003050755.1; 22 scaffolds), the genome of strain YX-36 was re-sequenced by the Department of Energy Joint Genome Institute (JGI) using the PacBio Sequel II instrument. The updated assembly, available at NCBI (NCBI RefSeq assembly GCF_041080815.1) is complete, which facilitated the mapping of the transposon mutants. The mapping analysis of the magic pool transposon library allowed us to identify what genetic ‘parts’[37] were more effective for whole-genome transposon mutagenesis in strain YX-36. The results showed that for part2 (the promoter for the drug resistance CDS) and part3 (the drug resistance CDS), the part2.7 and part3.3 were the most effective for generating transposon mutants (see section 2.2; **Fig. 2A**). For part5 (promoter for the transposase), the promoters driving the *mariner* transposase gave ∼25X more mapped reads than the Tn5 transposase. Several specific part5’s seemed to work more or less similarly including part5.18 (**Fig. 2A**). Based on these data, we decided to use the single reassembled vector pTGG43_NN2 (in *E. coli* WM3064, strain AMD691) that was previously used to make RB-TnSeq libraries in the Bacteroidota species *Echinicola vietnamensis*, *Pedobacter* sp. and *Pontibacter actiniarum*[37]. This vector is a combination of some the identified optimal parts for YX-36: part1 (mariner transposase), part2.7 (*dnaE* promoter from *Bacteroides thetaiotaomicron*), part3.3 (*ermBP* Erm resistance gene from pHLL24[64]), part4 (barcoded backbone) and part5 mar_p5.18 (Dxs promoter from *Belliella baltica*).

**Figure 2.**
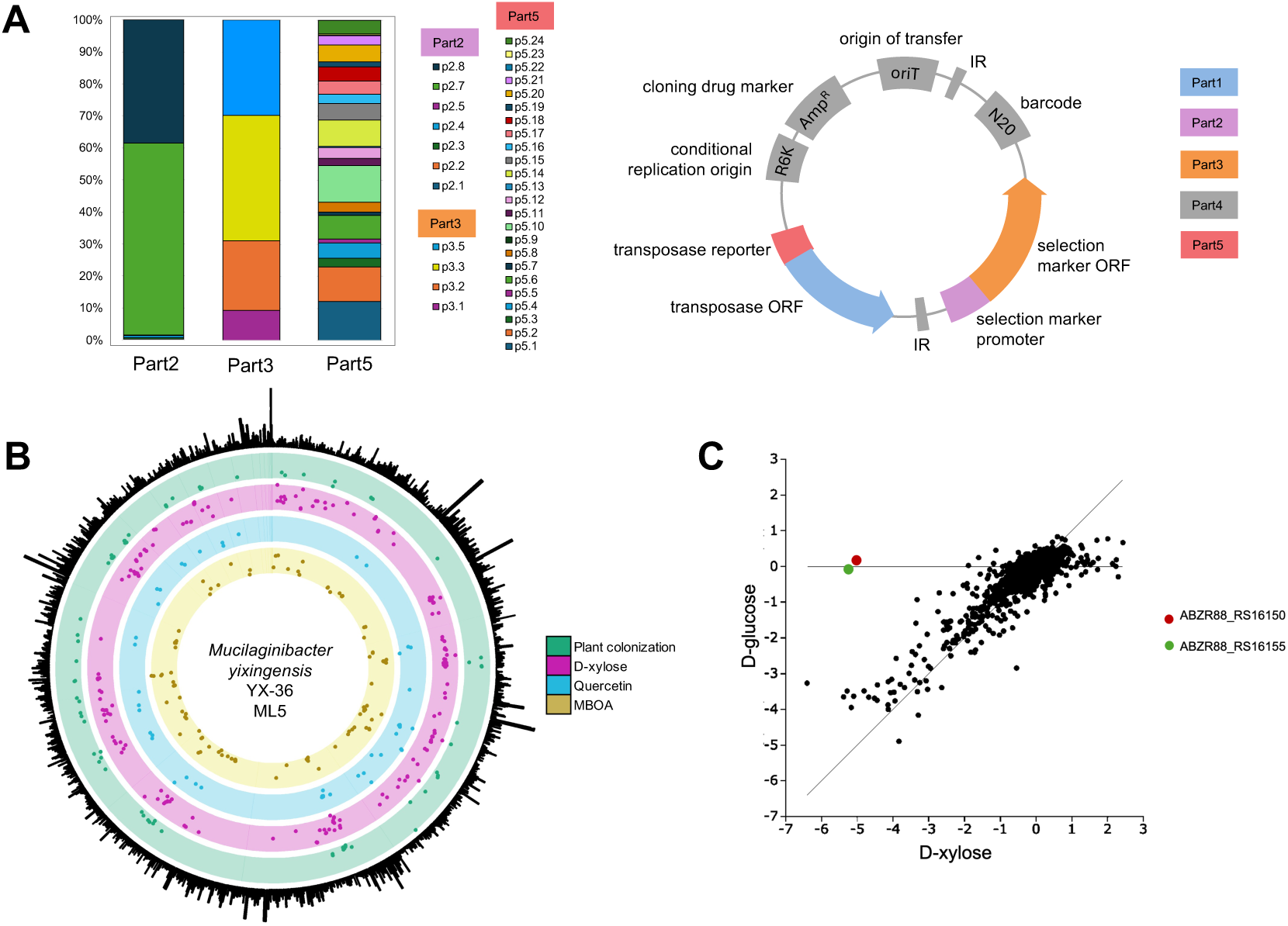
**A**) Preference of *Mucilaginibacter yixingensis* YX-36 for different vector parts in the Erm *mariner* magic pool. The fraction of each of part2, part3, and part5 variants in the preliminary mutant library for YX-36 as determined by DNA barcodes identified by TnSeq is shown. DNA part sequences and sources are available from Liu et al.[37]. The basic structure of the transposon delivery vector with the different parts indicated in colors is indicated; figure adapted from[37, 50]. **B**) Genome-wide map of YX-36 fitness genes. From outside to inside: total reads per gene from TnSeq mapping data, genes involved in plant colonization 7 days post inoculation (green), during growth in defined minimal media with D-xylose as the carbon source (pink), in stress tolerance to quercetin (blue) and 6-methoxy-2(3H)-benzoxazolone (MBOA; yellow). **C**) Correlation between averaged fitness data for D-xylose and D-glucose. ABZR88_RS16155 (coding for xylulokinase) and ABZR88_RS16150 (encoding for xylose isomerase) are shown.

Our final RB-TnSeq *mariner* library was named *Mucilaginibacter*_YX36_ML5. Subsequent TnSeq resulted in a pool of 61,393 mutant strains with mapped insertions and unique barcodes at 56,102 different locations distributed across the genome (**Fig. 2B**) at an average rate of 1 insertion location every 95 bp. From 4,390 protein-coding genes in the genome of YX-36, we found central mutations in 3,799 genes (86.5% of the genome). We collected fitness data for plant colonization, carbon metabolism, motility, temperature, and stress tolerance (**Table S2**) for 3,566 genes. The remaining genes do not have gene fitness data because they have few or no mutants in the library (many are likely essential genes) or mutants in these genes had a low number of reads in the Time0 samples (**Table S3**).

For each experiment that we conducted with *Mucilaginibacter*_YX36_ML5, gene fitness values were calculated as the log2 of the ratio of relative barcode abundance following library growth in a given condition (i.e. plant colonization) divided by relative abundance in the Time0 sample. A negative fitness (fitness ≤ 1) means that the mutants in that gene were depleted in that given condition; these are called ‘important’ genes. A positive fitness (fitness ≥ 1) means that the mutants in that gene are enriched in that condition relative to the typical mutant in the library; these are called ‘detrimental’ genes.

### 3.3 Validation of the *Mucilaginibacter*_YX36_ML5 mutant library

After generation of *Mucilaginibacter*_YX36_ML5, we used it with single monosaccharide compounds with the aim to validate the constructed library. The results (**Table S2**) obtained show that our mutant library is accurate, which validates it as a promising tool for gene function studies in environment-related *Mucilaginibacter* species. Specifically, the adjacent genes ABZR88_RS16155 (coding for xylulokinase) and ABZR88_RS16150 (encoding for xylose isomerase) are important for metabolism of D-xylose (**Fig. 2C**), which is the main residue in plant xylans[4], but not of other carbon sources we profiled including D-glucose. The role of these genes in xylose metabolism is well characterized in model organisms[65]. Interestingly, ABZR88_RS16150 is also important for colonization of plants (see section 3.4.4; **Table S2**). Another example of such validation is ABZR88_RS03755 (coding for a carbohydrate kinase/fructokinase, the limiting enzyme in fructose metabolism), which is only needed for growth in our experiments with D-fructose as the sole carbon source.

### 3.4 Identification of genes contributing to fitness in strain YX-36 across diverse *in vitro* conditions

In order to gain insight into different lifestyles and abilities related to environmental Bacteroidota[22, 52], we tested the YX-36 mutant library across several conditions (**Table S2**), such as metabolism of different carbon sources, gliding motility, and stress tolerance to different compounds.

#### 3.4.1 Metabolism of carbon sources

Bacteroidota are mainly able to orchestrate the degradation of complex glycans by organizing the key players into PULs, which allow high level production of specific CAZymes only when the corresponding substrate is abundant[4, 25]. PULs are sets of co-localized genes that usually encode a SusCD pair: a TonB-dependent transporter (SusC) that works with a glycan-capturing lipoprotein (SusD). In Bacteroidota glycan-degrading systems, secreted CAZymes partially depolymerize the polysaccharide to oligosaccharides which are imported into the periplasm by transporters encoded by a *susCD* gene pair, and subsequently degraded in the periplasm by other sugar-cleaving enzymes, away from competing organisms[25].

One powerful tool to identify PULs in Bacteroidota strains is the PULDB[60], which displays predicted PULs and CAZyme clusters (devoid of SusCD). Among the >40 *Mucilaginibacter* genomes in PULDB, the number of predicted PULs ranges from a dozen in *M. ginkgonis* HMF7856 and *M. ginsenosidivorans* Gsoil 3017, to almost a hundred in *M. sabulilitoris* SNA2 and *M. paludis* DSM 18603. In comparison, 45 PULs are predicted in *M. yixingensis* YX-36 (**Fig. S2**, **Table S4**). When the specificities of CAZyme (sub)families gather in a PUL match known glycan structure, it allows to predict the PUL substrates and likely growth ability of the species. For example, in *M. yixingensis* YX-36, predicted PULs 22, 32, and 44 likely target xylan, ⍺-1,4/6-glucans (e.g. starch, pullulan), and β-1,3-glucans (e.g. plant mixed-β-1,3/4-glucans or fungal β-1,3-glucans) respectively. Additional PULs are predicted to contain CAZymes that target complex and intertwined pectin polysaccharides.

Expert CAZy annotation of the improved YX-36 genome assembly shows a high number of enzymes belonging to different families (**Table S4**). More precisely, we found 95 glycosyltransferases (GT), which form glycosidic bonds; 35 carbohydrate esterases (CE), that hydrolyze carbohydrate esters; 16 carbohydrate-binding modules (CBM), that adhere to carbohydrates; 202 glycoside hydrolases (GH), that hydrolyze and/or rearrange glycosidic bonds; and 32 polysaccharide lyases (PL), which perform non-hydrolytic cleavage of glycosidic bonds. The number of PLs and CEs in *M. yixingensis* is the highest among the *Mucilaginibacter* species currently annotated (**Fig. 3**, **Fig. S3**). Our computational analysis agrees with previous findings on high numbers of CAZymes in the genomes of *Mucilaginibacter* species, especially GHs[9]. While CAZymes can also be secreted by the phylum-specific T9SS (**Fig. 4A**)[66], in strain YX-36 we only found two PLs that display a C-terminal domain (CTD) indicating their secretion by the T9SS.

**Figure 3.**
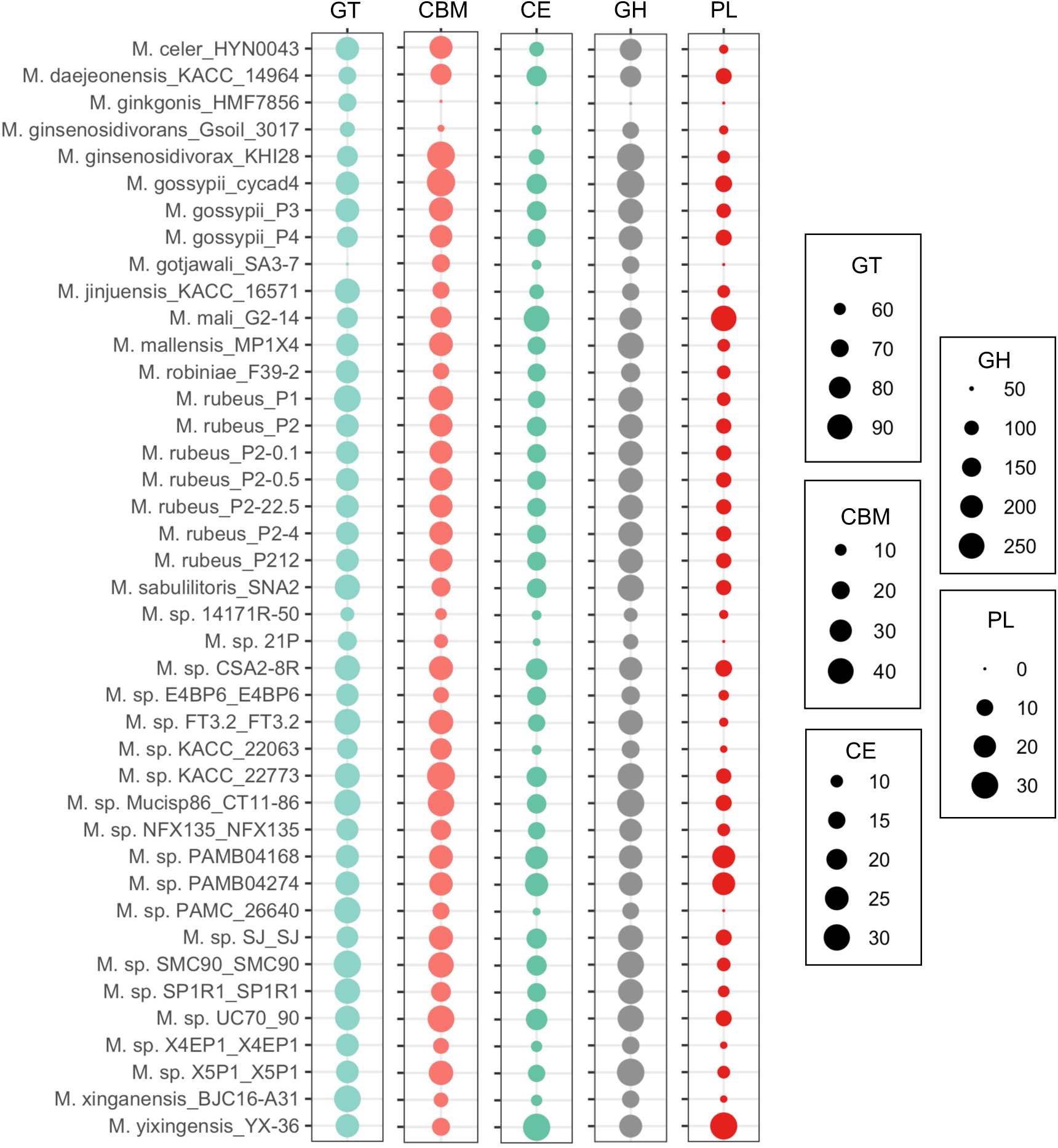
Total number of CAZymes among the 41 *Mucilaginibacter* strains in the CAZyme database. GH, glycosyl hydrolase; GT, glycosyl transferase; PL, polysaccharide lyase; CE, carbohydrate esterase; CBM, carbohydrate binding module.

**Figure 4.**
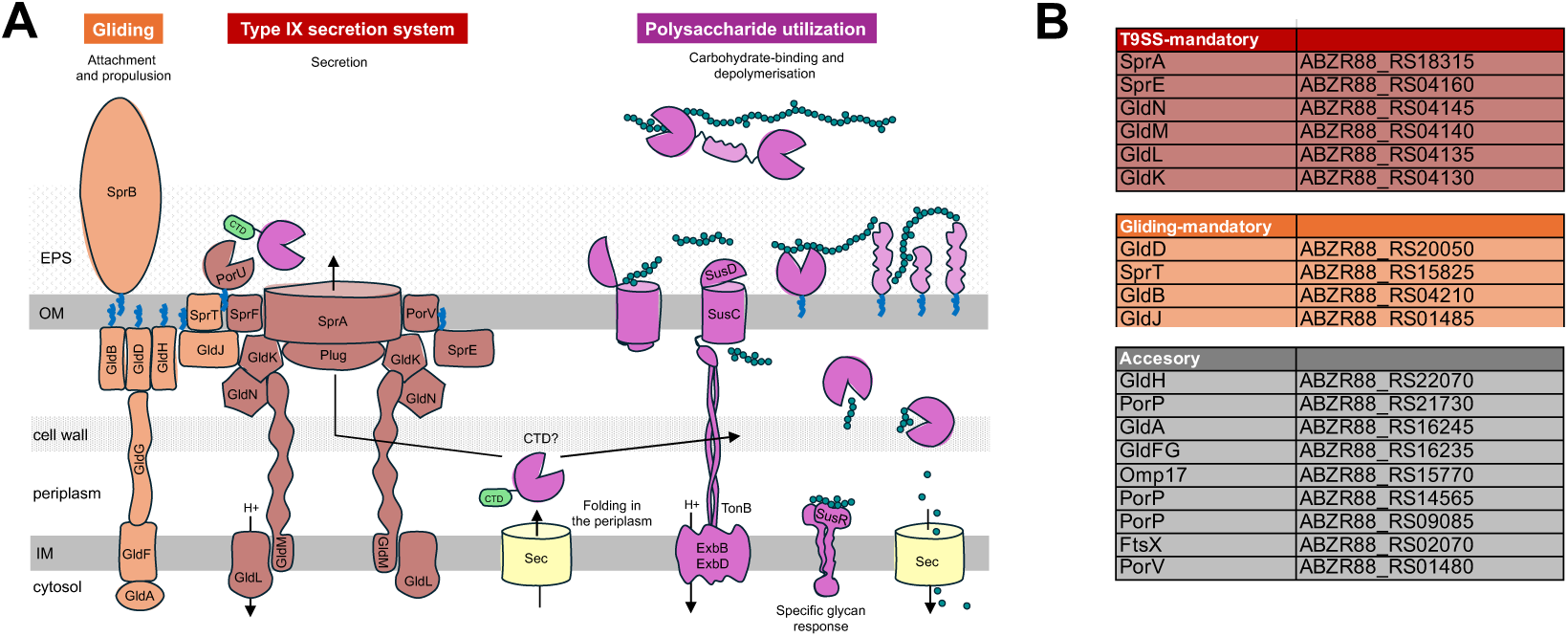
**A**) Overview of the connection between polysaccharide utilization loci (PULs; in pink) and the ability to glide rapidly over solid surfaces (in orange) by the type IX secretion system (T9SS, in red). EPS, exopolysaccharide; IM, inner membrane; OM, outer membrane. Blue wavy lines on proteins symbolize lipid anchors for lipoproteins. Figure modified from Larsbrink and McKee[4]. **B**) T9SS and gliding mandatory components in *Mucilaginibacter yixingensis* YX-36, as assessed with the T9GPred tool[72].

To represent the diversity of compounds degraded by environmental Bacteroidota, we selected different polysaccharides, disaccharides, and monosaccharides. For plant-related polysaccharides we used ⍺-1,4-glucans (e.g. starch, polygalacturonic acid) and β-1,4-glucans (e.g. carboxymethylcellulose). As disaccharides we used derivatives of the above polysaccharides; including D-cellobiose (a subproduct of carboxymethylcellulose) or D-maltose (produced after the hydrolysis of starch). As monosaccharides we used some of the basic units of the above, such as D-glucose, L-arabinose, and L-rhamnose. From the 14 carbon sources that we profiled, we identified genes specifically important for the catabolism of each, as specific phenotypes can be used to more readily assign gene function compared to pleiotropic genes[56]. One example is the cluster ABZR88_RS14505-ABZR88_RS14520 which is important for fitness on L-arabinose, with three genes associated with decreased fitness: a ribulokinase adjacent to an L-ribulose-5-phosphate 4-epimerase (ABZR88_RS14505-ABZR88_RS14510) and an L-arabinose isomerase (ABZR88_RS14520). These three genes correspond to the arabinose operon first described in *E. coli*[67]. Within this cluster, a predicted GH27 enzyme (ABZR88_RS14515) had no clear impact on fitness, despite more than 50% identity to β-L-arabinofuranosidases. One possible explanation may be the presence of a redundant gene in the YX-36 genome (i.e. ABZR88_RS12975, 31% homology), which would mask the phenotype due to compensation. Other cases are ABZR88_RS03755 and ABZR88_RS13180-ABZR88_RS13190, involved in fructose metabolism and D-cellobiose metabolism, respectively.

Of the four polysaccharides we profiled, we could link two to at least one PUL because multiple genes in the PUL were important for fitness on that substrate. First, we found that PUL32 was mildly important for growth on alginate, with 5 genes (ABZR88_RS13305:AZBR88_RS13325) with significant phenotypes. In contrast to the single PUL we identified for alginate, we found that 4 different PULs were important for growth on polygalacturonic acid (PUL3, PU14, PUL 19, PUL20), although not all predicted genes within each PUL were required for fitness. In PUL3, the SusCD pair was not important, but other genes including ABZR88_RS02125 (pectinesterase family protein) had mild but significant phenotypes. In PUL14, only the *susCD* genes (ABZR88_RS06240-ABZR88_RS06245) were important for growth on polygalacturonic acid, while the other genes were dispensable. While PUL19 is predicted to encode two separate SusCD pairs, only the ABZR88_RS08375: ABZR88_RS08380 system was important for fitness. Lastly, in PUL20, 3 of the 5 predicted genes including the SusCD homologs (ABZR88_RS08490:ABZR88_RS08500) were required for optimal growth on the polysaccharide. Our results demonstrate that in certain instances, components of multiple PULs can contribute to the catabolism of a complex polysaccharide.

Although our main focus was genes with a negative fitness, we also analyzed detrimental genes (i.e. with positive fitness, meaning that the mutation in that gene is beneficial and causes a boost in fitness). We found that a cluster (ABZR88_RS07190-ABZR88_RS07205 within PUL16, **Table S4**) coding for two pairs of GH2 and GH3 enzymes is detrimental (fitness ≥ 1) for metabolism of D-melibiose, a disaccharide present in plants. While melibiose breakdown requires an ⍺-galactosidase, these four enzymes are β-glycosidases, which close homologs rather suggest the degradation of polysaccharides made of β-galactose, β-glucose, and β-glucuronic acid, such as in soil rhizobial exopolysaccharides[68]. This phenotype is exclusive to transposon mutations inserting on the positive strand of these genes, and the probable overexpression of the downstream gene ABZR88_RS07210 (coding for an alpha-galactosidase). In support of this, ABZR88_RS07210 is mildly important for melibiose utilization, and thus overexpression of this gene would provide a fitness benefit on this substrate. This finding very closely mirrors what was previously observed in *Bacteroides thetaiotaomicron* in mice, where transposon mutants driving overexpression of a gene important for melibiose utilization have a large competitive advantage[41]. Another example is a cluster (ABZR88_RS04740-ABZR88_RS04750) that is detrimental for metabolism of D-xylose, D-cellobiose, D-xylose, D-arabinose, polygalacturonic acid, and gliding. One of these genes belongs to family GT2, questioning the possible importance of the glycosylation of gliding proteins similarly as for motility by flagella in Gram-negative[69]. In total, we identified genes involved in carbon metabolism (|fitness ≤ 1|) in 15 out of the 45 different PULs in strain YX-36 (**Table S2**, **Table S4**). In the future, similar experiments with additional plant complex glycans could provide insightful information to understand gene function and reannotate hypothetical genes in strain YX-36.

#### 3.4.2 Gliding motility

T9SS is tightly linked to gliding motility through the involvement of the T9SS in adhesin export and possibly through the use of a shared motor complex[60, 66, 70]. The ability to glide, movement on surfaces without the aid of external appendages such as flagella, is widespread among Bacteroidota and it allows an active search for nutrients[4, 70] as well as rapid secretion and release of CAZymes into the environment[66].

The central element of the T9SS is SprA, which forms a 36-strand single polypeptide transmembrane β-barrel. T9SS substrates are large (100-650 kDa) multi-domain proteins which fold in the periplasm before being exported[71]. SprA was recently identified as the largest single polypeptide transmembrane β-barrel identified to date, far exceeding the size of the 26-strand lipopolysaccharide transporter LptD[71]. Some other components of the T9SS are GldKLMN and PorQUVZ (**Fig. 4A**). The exact function of many of the T9SS components are unclear. In *M. yixingensis* YX-36 there are more than 100 genes annotated as related to gliding and T9SS. The gliding (GldBDJ, SprT) and T9SS (GldKLMN, SprAE) mandatory components are present in this strain, as we assessed with the T9GPred[72] tool (**Fig. 4B**), meaning that YX-36 has T9SS and is able to glide.

We confirmed that the predicted mandatory gliding genes GldBD, SprT (ABZR88_RS20050, ABZR88_RS15825, ABZR88_RS04210 (**Fig. 4B**) were important (fitness ≤ 1) for gliding motility in strain YX-36 (**Table S2**). Gene GldJ (ABZR88_RS01485) was not identified as involved in gliding in our screening, but its homologue ABZR88_RS04130 was. Other expected genes that we found are ABZR88_RS01480 (PorV), ABZR88_RS04135 (GldJ homologue), ABZR88_RS04135 (GldL), ABZR88_RS04140 (GldM), ABZR88_RS04145 (GldN), ABZR88_RS16235 (GldG) and ABZR88_RS18315 (cell surface protein SprA) (**Table S2**). Many of the above genes have high positive cofitness, and their homologues have previously been related to gliding motility in Bacteroidota[73]. We also identified novel genes related to gliding motility: ABZR88_RS02055 (signal peptidase I), ABZR88_RS20165 (ribosome silencing factor), and ABZR88_RS00640 (serine hydroxymethyltransferase), the latter being important for growth in multiple other conditions as well including plant colonization (**Table S2**).

#### 3.4.3 Stress tolerance

*Mucilaginibacter* species are resistant to diverse stresses, such as cold temperatures, metal salts, and antibiotics[22, 55, 74]. One possible explanation of this tolerance is the abundant exopolysaccharides that they synthesize[75]. We assayed *Mucilaginibacter*_YX36_ML5 in different stress conditions: temperature (i.e. 15°C, 21°C, 30°C, 37°C), metal salts (i.e. NaCl, CuCl2, KF, K2HPO4), antibiotics (i.e. kanamycin), and plant-related molecules (i.e. 6-methoxy-2(3H)-benzoxazolone or MBOA, quercetin, apigenin, esculin).

From these data, we found that ABZR88_RS11280 (TonB-dependent receptor) was important (fitness ≤ 1) for tolerance to quercetin (and mildly important for gliding motility), but not the adjacent TonB-dependent receptor (ABZR88_RS11275). We also found that ABZR88_RS11640 (sodium-translocating pyrophosphatase), ABZR88_RS22105 (voltage-gated chloride channel family protein), and ABZR88_RS14150 (magnesium transporter) were needed for growth in presence of KF. Homologs of both of these genes were also found to be required for fluoride tolerance in *Pedobacter* sp.GW460-11-11-14-LB5[56]. We also identified phenotypes for several genes coding for RND (resistance-nodulation-cell division) family transporters[76], such as the efflux RND transporter periplasmic adaptor and permease pair ABZR88_RS03435-ABZR88_RS03440, which was detrimental for stress tolerance to CuCl2 (**Table S2**).

Interestingly, we identified that several components of the Bacteroidota-specific T9SS, namely ABZR88_RS01480 (T9SS outer membrane channel protein PorV), ABZR88_RS18315 (cell surface protein SprA), ABZR88_RS04140 (gliding motility protein GldM), ABZR88_RS04145 (GldN), ABZR88_RS04210 (GldB), ABZR88_RS20050 (GldD), and ABZR88_RS16235 (GldG) are needed for tolerance to the plant-associated stressors MBOA, quercetin, apigenin, and high K2HPO4 (**Table S2**). Many of these genes have high positive cofitness, which is the correlation between the fitness patterns of two genes. Genes with high cofitness tend to perform shared functions.

As other Bacteroidota, members of the *Mucilaginibacter* genus are also very resistant to different antibiotics. Multiple antimicrobial resistance pathways have been characterized in *Mucilaginibacter* species[77]. Strain YX-36 is resistant to kanamycin, among others. Interestingly, it was recently described that strain *Mucilaginibacter* 21p has cross-resistance mechanisms that confer resistance to both antibiotic and metal salts[55]. In agreement with this, we observed that genes ABZR88_RS12790 (glycosyltransferase), ABZR88_RS12785 (flippase), ABZR88_RS17545 (glycosyltransferase GT2 family), ABZR88_RS09925 (ABC transporter ATP-binding protein), ABZR88_RS09930 (ABC transporter permease), ABZR88_RS12795 (UDP-galactopyranose mutase), and ABZR88_RS17535 (oligosaccharide flippase family protein) are all important for stress tolerance to the salts KF and NaCl and the antibiotic kanamycin, but not for other stressors such as CuCl2, apigenin, quercetin or MBOA (**Table S2**). These genes are probably involved in the import of components for the production and externalisation of a polysaccharide biofilm key to bacterial protection and nutrient storage.

### 3.5 Identification of plant colonization genes in strain YX-36

*Mucilaginibacter* species have been isolated from the root and rhizosphere of different plants, such as *Brachypodium*, switchgrass, and rice[49, 53, 78, 79]. Some strains have been found to be positively correlated with crop biomass and yield[80]. Others are increased in abundance and proportion within a community when the amount of exuded compounds increases[81]. Recently, we described a *Mucilaginibacter* strain that could efficiently colonize the rhizosphere of *B. distachyon* plants in competition with other bacterial species[48]. Although numerous studies focus on the identification of plant colonization fitness genes in bacteria belonging to the phylum Pseudomonadota[36], few or no data is available on plant-related Bacteroidota. With the aim to bridge this gap and identify plant colonization strategies in environmental Bacteroidota, we evaluated the fitness of *Mucilaginibacter*_YX36_ML5 in the rhizosphere of *B. distachyon* plants at 7 and 21 dpi (**Fig. 5A**). This was made to consider the dynamism of the colonization process[41]. On a global scale, our data indicate that a greater number of genes are specifically implicated in plant colonization at 21 dpi compared to 7 dpi (**Table S2**), potentially reflecting a more stringent adaptive response to the rhizosphere environment or altered nutrient availability. Nonetheless, additional investigations incorporating more temporally distant sampling points are warranted to substantiate these observations.

**Figure 5.**
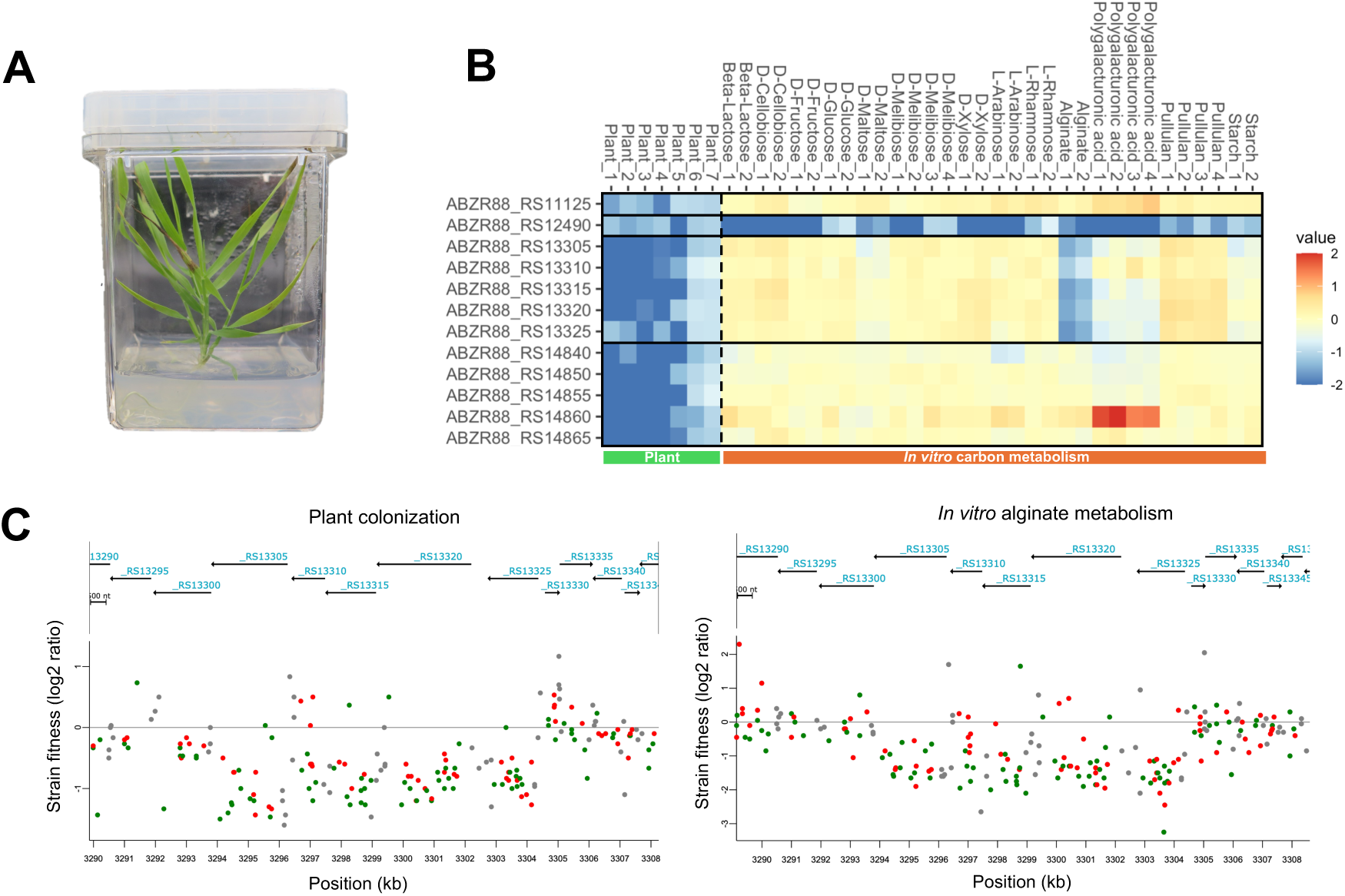
**A**) Overview of *Brachypodium distachyon* Bd21-3 plants **B**) Heatmap of fitness values for a selection of plant rhizosphere colonization genes across different plant and *in vitro* tests. Gene fitness values are bounded from -2 to 2. **C**) Strain fitness for selected genes (from ABZR88_RS13290 to ABZR88_RS13345) across average plant assays and alginate metabolism. Average of replicates is shown. Strains in the central 10-90% of a gene are color coded (green or red dots) by the insertion’s strand. Strains near the edge of a gene are not shown as being associated with that gene (grey dots).

Several of the genes that we found to be involved (|fitness| ≥ 1) in plant colonization by strain YX-36 are related to amino acid synthesis. Examples are ABZR88_RS06865 (imidazole glycerol phosphate synthase subunit HisH), ABZR88_RS06870 (imidazole glycerol-phosphate dehydratase HisB), ABZR88_RS11470 (glutamine synthetase), and ABZR88_RS11615 (threonine ammonia-lyase IlvA) (**Table S2**). Orthologues of these genes have previously been associated with plant colonization in phylogenetically distant species[36, 82, 83], which shows that amino acid synthesis is required by different bacteria for efficient plant colonization[36]. Genes within other functional categories that we find involved (|fitness| ≥ 1) in rhizosphere survival are: ABZR88_RS00290 (mannonate dehydratase), ABZR88_RS16205 (bifunctional 4-hydroxy-2-oxoglutarate aldolase), ABZR88_RS12770-ABZR88_RS12800 (lipopolysaccharide biosynthesis proteins), and ABZR88_RS04145 (gliding protein GldN). They correspond to phenotypes commonly found in plant-associated bacteria, such as biofilm formation and motility[84, 85].

Interestingly, our findings reveal that largely phylum-specific genes (e.g. PULs, CAZymes) are needed for fitness in the plant niche, allowing Bacteroidota strains privileged access to nutrients in the rhizosphere that remain inaccessible to other phyla like Pseudomonadota. Examples are the genes ABZR88_RS02185-ABZR88_RS02190 (pectinesterase family proteins, CAZyme family CE8, in PUL4), ABZR88_RS11125 (glycoside hydrolase, CAZyme family GH88, in PUL23), and ABZR88_RS12490 (⍺-1,4-glucan branching protein GlgB, displaying families CBM48 and GH13 domains, and related to ⍺-glucan metabolism), the later showing negative fitness in the presence of multiple carbon sources including D-xylose, D-fructose, and L-arabinose. Another example of a plant-associated phylum-specific cluster is ABZR88_RS13305-ABZR88_RS13325, coding for a glycoside hydrolase GH31 family protein, a SusE domain-containing protein, a RagB/SusD family nutrient uptake outer membrane protein, a TonB-dependent receptor and a DUF6377 domain-containing protein (**Fig. 5B**, **Table S4**). These genes, all belonging to PUL32, are needed for both *Brachypodium* colonization and alginate metabolism (**Fig. 5B**, **Fig. 5C**). Several genes within PUL2, PUL4, PUL7, PUL16, and PUL17 also were needed for colonization of the plant niche (**Table S2, Table S4**). Finally, we also identified a group of genes belonging to PUL39 that were mildly but significantly important for plant colonization: ABZR88_RS14840 (sodium/sugar symporter), ABZR88_RS14850 (RagB/SusD family nutrient uptake outer membrane protein), ABZR88_RS14855 (TonB-dependent receptor), ABZR88_RS14860 (inositol oxygenase family protein) and ABZR88_RS14865 (LacI family DNA-binding transcriptional regulator) (**Fig. 5B**, **Table S2**). The specific substrate(s) recognized by PUL39 are not clear, as we did not detect a phenotype for these genes in our *in vitro* dataset, and close homologs of these proteins have not yet been characterized. Our results demonstrate that efficient plant colonization in YX-36 requires the activity of multiple PULs, providing evidence that proteins characteristic of Bacteroidota are important for this process, likely mediated via the degradation of polysaccharides and their break-down products. Future experiments could focus on deeper characterization of the identified PULs important for *Brachypodium* colonization, as well as including more diverse plant species to determine if these colonization genes are specific to certain hosts.

## 4. Conclusion

Bacteroidota species are trending in microbiology research due to their exceptional adaptive capacity to proliferate in diverse environments, but the genetic determinants underlying the various lifestyles are not yet clear. In spite of the interest raised in the last years in the environmental members of this phylum, it remains a challenge to study the pathways and mechanisms related to fitness in plant hosts. This is in part due to the high intrinsic antimicrobial resistance of Bacteroidota and therefore their challenging genetic manipulation, as genome-wide mutant-based screening methods (e.g. RB-TnSeq) are usually based on selection methods which rely on antibiotic resistance.

In this study we attempted to construct barcoded mutant libraries in a collection of environment-related Bacteroidota species and succeeded in generating one in *M. yixingensis* YX-36. We produced fitness data for YX-36 in several *in vitro* and plant assays. Our data confirms that RB-TnSeq is a promising tool to unravel novel plant colonization genes in non-Pseudomonadota. Notably, our results highlight that T9SS-related genes and largely phylum-specific genes (e.g. PULs) are important for plant colonization, and may give these Bacteroidota strains unique access to nutrients or niches that other phyla (e.g. Pseudomonadota) cannot access. We present here strong candidates for continued functional characterization through controlled *in vitro* and *in vivo* experiments, and present an exciting avenue for further research. Deciphering the poorly explored functional potential of plant-associated Bacteroidota will contribute to the development of novel biotechnological, medical, and environmental applications of this understudied bacterial phylum. We hope that our mutant library will be an exciting resource for the community doing research in the poorly characterized genus *Mucilaginibacter*, and that our results will draw more attention to environmental Bacteroidota, especially those living in close association with plants and food crops.

## 5. Acknowledgments

The authors thank Morgan Price for processing fitness data, Kateryna Zhalnina for providing environmental Bacteroidota isolates, John Vogel and Mingqin Shao for providing Bd21-3 seeds, and the QB3 Genomics Center at UC Berkeley for sequencing.

## 8. Contributions

MT and AMD designed the study. MT constructed the YX-36 barcoded mutant library and performed fitness assays. AMD obtained funding. NT and VL performed CAZyme annotation and analyses. MT analyzed data, prepared figures and tables, and drafted the manuscript. MT, NT, VL and AMD wrote the final version of the manuscript.

## 10. Supplementary material

**Figure S1.**
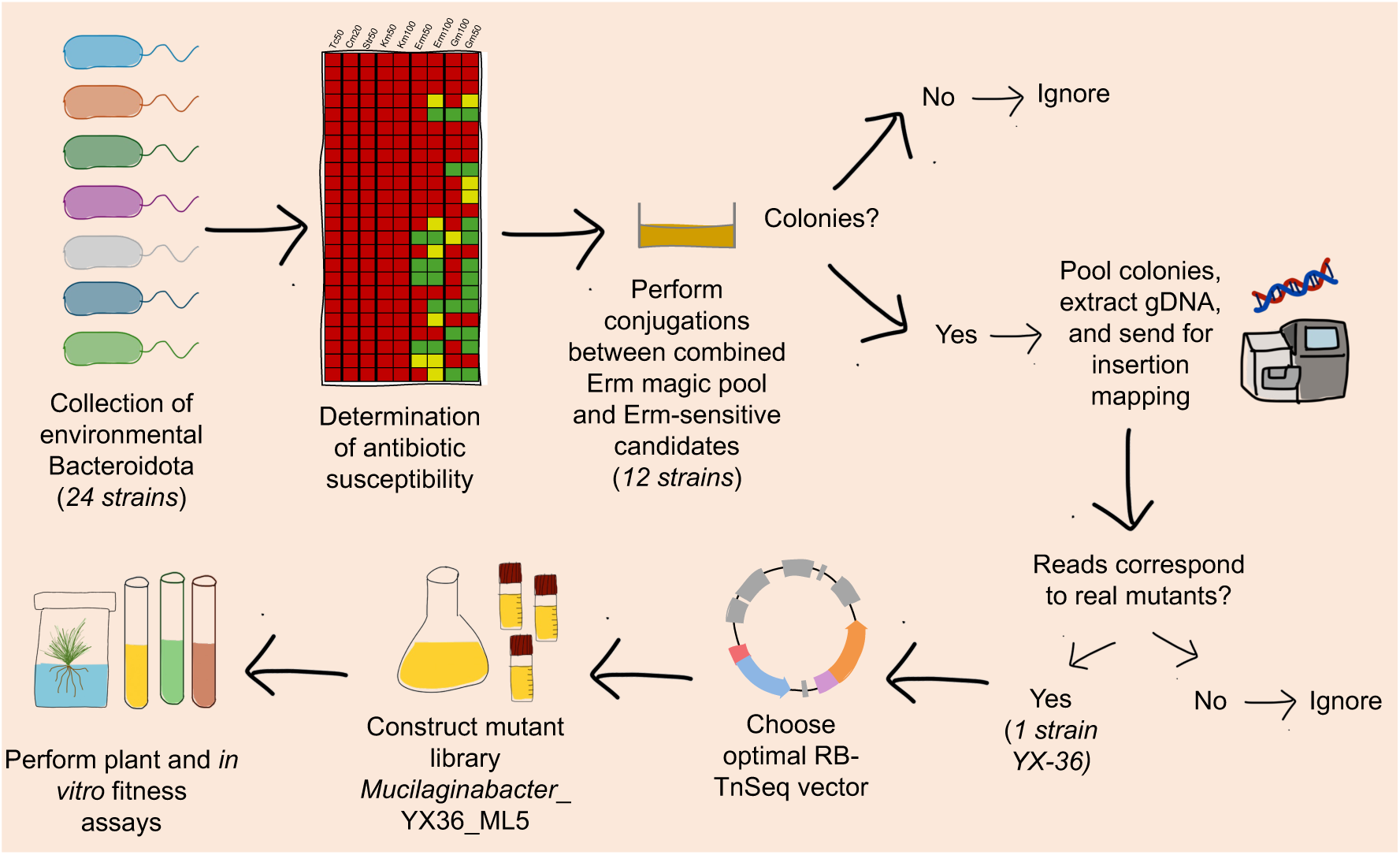
**O**verview of the methodology and the steps followed in this study to construct a mutant library in *Mucilaginibacter yixingensis* YX-36.

**Figure S2.**
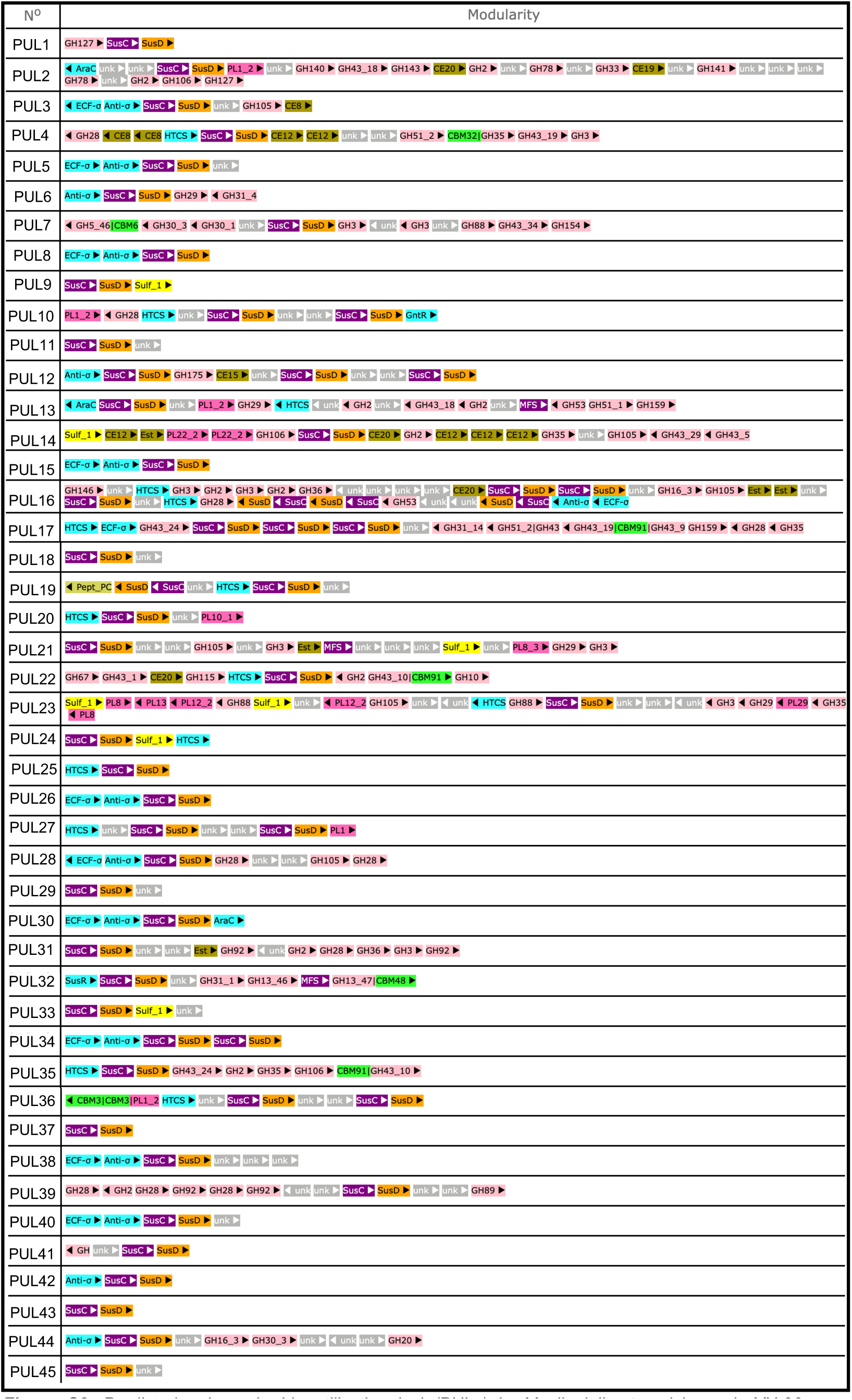
Predicted polysaccharide utilization loci (PULs) in *Mucilaginibacter yixingensis* YX-36 as determined by the PUL database (PULDB)[60].

**Figure S3.**
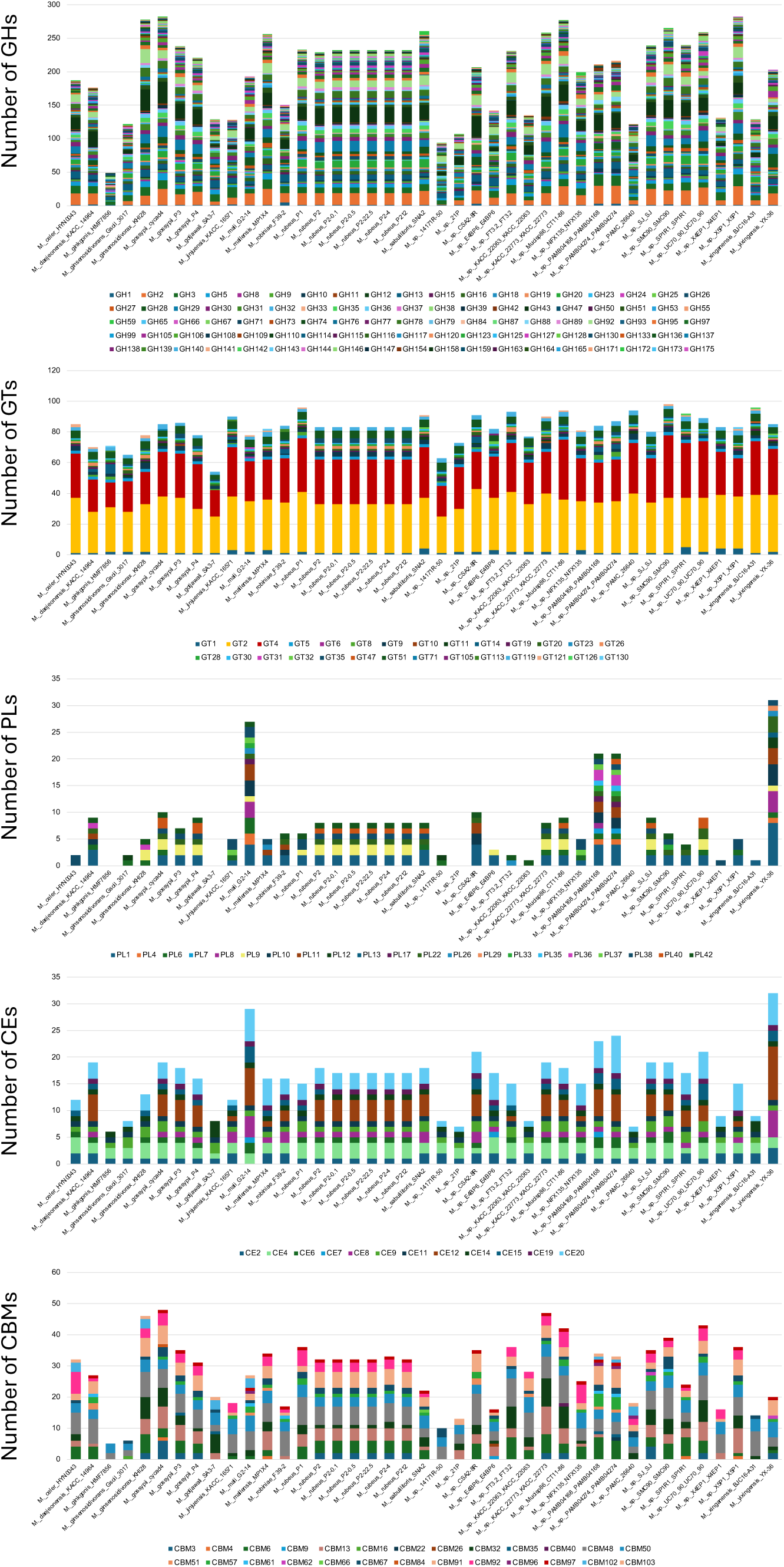
Distribution of CAZymes among the 41 *Mucilaginibacter* strains in the CAZyme database[26]. GH, glycosyl hydrolase; GT, glycosyl transferase; PL, polysaccharide lyase; CE, carbohydrate esterase; CBM, carbohydrate binding module.

**Table S1.** Summary of fitness assays conducted with *Mucilaginibacter*_YX-36_ML5.

**Table S2.** Fitness data of *Mucilaginibacter yixingensis* YX-36. Fitness values and t scores are shown. Assays publicly available at the Fitness Browser website (https://fit.genomics.lbl.gov) are noted.

**Table S3.** *Mucilaginibacter*_YX-36_ML5 gene information, including scaffold, begin, end, strand, GC content of the gene’s sequence and nTA (number of TA dinucleotides in the gene’s sequence; the mariner transposase usually inserts at TA).

**Table S4.** Predicted PULs and CAZymes in the genome of *Mucilaginibacter yixingensis* YX-36.

